# Simple fence modification increases floodplain land movement prospects for freshwater turtles

**DOI:** 10.1101/2020.12.03.409607

**Authors:** Nathan J. Waltham, Jason Schaffer, Justin Perry, Sophie Walker, Eric Nordberg

## Abstract

Feral pigs predate on freshwater turtles and damage wetland habitats in the process. Installing fences successfully averts access and damage, however, they become a barrier for freshwater turtles requiring land access during migration. We collected 161 turtles (*Chelodina rugosa*, *Emydura subglobosa worrelli, Myuchelys latisternum*) from twenty floodplain and riverine wetlands during post-wet (June-August) and late-dry season (November-December) surveys (2015-2018) in northern Australia. Wetlands were either fenced (150 × 150mm square, 1.05m high wire mesh) or not around the wet perimeter. Nine-seven percent of individuals caught in either fenced or unfenced wetlands had a shell carapace width greater than mesh width, of these 44 (46%) were captured inside fenced wetlands, while 50 were caught in unfenced wetlands. The remaining 35 were smaller than 150mm and would easily pass through fence mesh. Sixty-five turtles partook in a fencing manipulative experiment. Turtles with carapace widths wider than mesh often successfully escaped through fences by lifting one side of their shell and passing diagonally. In a second experiment where a piece of vertical wire (1500mmx300mm) was removed, turtles located gates after prospecting and trying to fit through meshing areas that were too small to pass through. Nine-two percent of turtles were able to locate and pass through gates, while 8% failed to locate a gate after 2 hours. Three turtles that did not use gates, and seemed to ‘give up’ and dug into the grass. Gates applied every 4m showed an 83% passage rate, every 2m was 91%, and while every 1m was 100%. Combing field and manipulative experiments revealed that large turtles will prospect and move along a fence until they find suitable passage. Applying turtle gates every 1–4m allows almost 100% passage, and if strategically applied in travel corridors, would minimize the need for large-scale clipping efforts around entire wetlands.

## 1. Introduction

Conservation fences are a way to ameliorate threatening processes from acting against individual species or for conservation of sensitive ecosystem habitats [1]. While conservation fences have been successful [2], they also have negative indirect effects on non-target species [3, 4], resulting in an ongoing conservation dilemma for managers [5, 6]. Emerging evidence suggests that fencing affects non-target species, for example, by disruption to dispersal processes, and increased mortality (via increased exposure to unfavourable conditions or predators; Spencer (7)). These impacts are greatest on vagile animals which have evolved behavioral life history traits that allow them to inhabit landscapes characterized by spatial and temporal variability, and are therefore susceptible to limited access to resources or responding to local pressures (predation, climate conditions). However, with every conservation fence there exists the opportunity to evaluate the design efficacy, and implement supplementary modifications and improvements as part of a process of continual improvement [3].

Wetlands (palustrine and lacustrine) located on floodplains away from riverine channels support rich aquatic plant and fauna communities [8]. During high water levels in flood, interconnecting riverine channels create a linking network of waterbodies that persist permanently or only in an ephemeral state [9, 10]. Aquatic organisms occupying wetlands face a shifting land-water margin, until connection is finally broken. This results in wetlands supporting a non-random assortment of aquatic and semi-aquatic species [11, 12]. The duration, timing and frequency that off-channel wetlands sustain lateral connection to primary rivers is a determining factor in broader aquatic ecology and production [13, 14]. In addition to connection, environmental conditions become important including water quality [15–17], access to shelter to escape predation, and available food resources [18]. Managers are increasing efforts to restore wetland ecosystem values [19], though access to data demonstrating success are limited, which becomes important when attempting to assess biodiversity return for the funding invested by government or private sector markets [20, 21].

Across northern Australia, feral pigs (*Sus scrofa*) contribute wide-scale negative impact on wetland vegetation assemblages, water quality, biological communities and wider ecological processes [22, 23]. Feral pigs have an omnivorous diet supported by plant roots, bulbs and other below-ground vegetation throughout terrestrial and wetland areas [24]. This feeding strategy has a direct negative impact on wetland aquatic vegetation [25, 26], which gives rise to soil erosion, benthic sediment resuspension, and reduced water clarity and eutrophication which is particularly critical late-dry season. Only a few studies have quantified the negative impacts feral pigs have on coastal wetlands [26–29], limiting the ability of land managers to measure the benefits of feral pig destruction [30], or indeed other large invasive species [31]. Strategies focused on reducing or removing feral pigs from the landscape have been employed since their introduction to Australia [30], including poison baiting, aerial shooting, and trapping using specially constructed mesh cages [32]. Attempts to exclude feral pigs have also included building exclusion fencing for conservation outcomes by directly limiting access to essential resources [33]. The installation of fences around wetlands has only recently been examined in Australia [26, 27], with results suggesting that fences may prevent non-target terrestrial fauna access which becomes particularly relevant late-dry season when wetlands are regional water points in the landscape. While small terrestrial species including birds, snakes and lizards can still access fenced wetlands [32], freshwater turtles movement may be hindered. To this end, the inherent problem of wildlife fencing needs further consideration [6] as part of broader wildlife conservation and resource management strategies.

Globally, freshwater turtles are at risk of extinction due to landscape changes including poor habitat quality, fragmentation or total habitat loss [23, 34], nest predation [7, 35], or changes in hydrology either through direct water extraction or regulation [36], and climate change [37]. In northern Australia, a number of freshwater turtle species inhabit seasonal wetland complexes [38] and will employ terrestrial locomotion to exploit ephemeral food supplies, to lay eggs or escape drought. Accessing terrestrial areas expose turtles to new hazards such as desiccation, and predation by other terrestrial fauna [39, 40]. Freshwater turtles hold important cultural values, which has led to funding feral control programs to install fences to protect turtles. The use of wetland perimeter fencing is now widespread in northern Australia, which has improved protection of aquatic vegetation and water quality [26]. However, fencing does still pose concerns relating to whether turtle movement is impeded.

As part of a broader feral pig abatement partnership between government, indigenous community, and research agencies [32], our aim here was to evaluate the potential effect that wetland exclusion fencing has on the population demographics of freshwater turtle species inhabiting floodplain and riverine wetland complexes in northern Australia. Specifically, we examined shell morphology in relation to fence dimension characteristics from turtle populations captured in fenced and unfenced wetlands to determine the proportion of individuals whose mobility across the landscape would be restricted because of fencing. Extending on the field observations and previous studies which have shown that turtles will persist in their attempts to overcome barriers to movement between wetlands [5], we tested simple ‘turtle gates’ on a commonly used exclusion fence to increase services provided by wetlands and mitigation efforts.

## 2. Methods

### 2.1 Description of study system

We studied freshwater turtles occupying floodplain and riverine wetlands between 2015 and 2018 within the Archer River catchment, Cape York Peninsula, Queensland (Figure 1). The headwaters rise in the McIlwraith range on the eastern side Cape York, where it flows and enters the western side of the Gulf of Carpentaria. The catchment area is 13,820km^2^, which includes approximately 4% (510km^2^) of wetland habitats, including estuarine mangroves, salt flats and saltmarshes, wet heath swamps, floodplain grass sedge, herb and tree *Melaleuca* spp. swamps, and riverine habitat. The lower catchment includes part of the Directory of Internationally Important Wetland network (i.e., nationally recognised status for conservation and cultural value) that extends along much of the eastern Gulf of Carpentaria, including the Archer Bay Aggregation, Northeast Karumba Plain Aggregation and Northern Holroyd Plain Aggregation. Two national parks are located within the catchment (KULLA (McIlwraith Range) National Park, and Oyala Thumotang National Park). Land use is predominately grazing.

**Fig 1.**
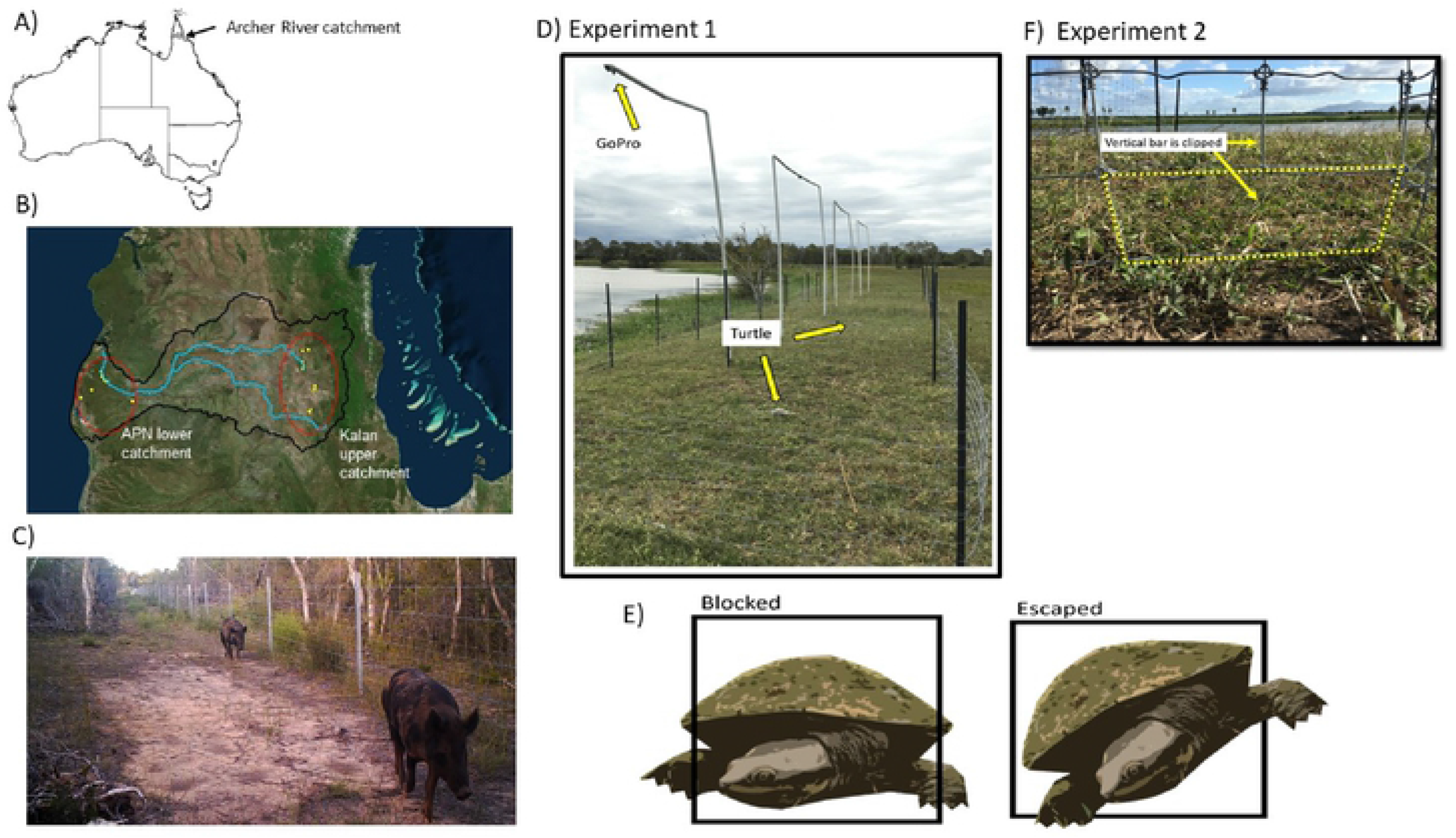
A) location of the Archer River catchment in northern Queensland, Australia; B) wetland sites on the coastal floodplain and mid catchment where feral pig fencing has been completed around wetlands preventing access (yellow circles); C) fenced wetland preventing pig access to coastal wetland (photo source S Jackson, Queensland Parks and Wildlife Services); D) Experiment I showing location of four arenas used to manipulate fencing gate design and turtle blockage/escaped; E) example of turtle blocked by fencing and escaped through fencing by angling body position; and F) Experiment 2 showing how gate opening was manipulated to test improvements to fencing design and allowing turtles to pass through fences.

Rainfall is tropical monsoonal, strongly seasonal with 90% of total annual rain occurring between November and February. Long term rainfall records for the catchment revealed highest wet season rainfall occurred in 1989/1999 (2515mm), while the lowest was 1960/1961 (563.5mm). Total antecedent rainfall for the wet season prior (Nov 2014 to Feb 2015) to this research was 1081mm, close to the 10^th^ percentile for historical records. The wet seasons experienced through the years prior to this study (2010 to 2015) were among the wettest on record, proximal to the 95^th^ percentile. The low rainfall experienced during this study may have contributed to short flood duration, and connection between wetlands and the Archer River.

Twenty wetlands were sampled including both floodplain and riverine wetlands that were not on the main flow channels, but rather on anabranches and flood channels that connect to the main channels only during high flow events (Waltham & Schaffer, in review). All wetlands in the region have been damaged by pigs (and cattle to a lesser extent) for the past 160 years [41, 42]. However, recently local indigenous community groups commenced a program of fencing wetlands to abate feral pig and cattle from accessing wetlands, in accordance with indigenous groups Kalan Enterprises, Aak Puul Ngangtam, and partners to meet the objectives of traditional owners [32].

### 2.2 Field methods – fenced and unfenced wetlands

Freshwater turtles were captured using specialized circular (820mm×2500mm) collapsible ‘cathedral-style’ traps [43] baited with canned sardines in vegetable oil. Generally, two traps were deployed in ~1.5m of water, spaced ~150m apart, mid-to-late afternoon (1500–1700hrs) and checked between 1000 and 1200hrs the following day. In some wetlands and at certain times of the year, low water levels rendered cathedral traps impractical. In these instances, turtles were passively sampled with unbaited fyke nets (1mm mesh, 0.5m height, single wing panel span 10m) set along the wetland margins. All traps were open and undisturbed overnight. Captured turtles were weighed, measured (following the morphometric codes in Table S1) and released back at the site of capture. In addition to trapping, the perimeters of fenced wetlands were searched for evidence of turtles either alive or dead trying to pilot through fences. If found, the morphometric data of turtles were recorded and added to the dataset.

### 2.3 Fencing manipulative experiment

#### Experiment 1 – fence mesh sizes

Four replicated field arenas were constructed on a flat grassy bank adjacent to a wetland lagoon near Townsville, Queensland (Figure 1D). Each arena (4×6×1m [L×W×H]) was constructed using 180cm star pickets to which we attached galvanized fencing (Southern Wire Griplock^®^ 80/90/15) identical to that used in feral pig management in the Archer River catchment. Fences were 90cm high and composed of 2.50mm wire with a standard 150mm gap between vertical strands. Eight horizontal strands of wire create 7 mesh panels which are arrayed in a vertically increasing graduated mesh design (mesh area [LxWmm] ‘large’ = 2316 ±81cm^3^; ‘small’ = 1540 ±46cm^3^) (Table S2). Generally, the smaller mesh size is used at the bottom of the fence to reinforce against the prospect of pigs digging under fences [32]. We tested the passage rates of turtles through these fences oriented with both the small (normal) and large (up-side-down) mesh panels at the bottom.

Sixty-five turtles (*Emydura macquarii kreftii*) were captured from waterbodies in close proximity to the experimental arenas. For every replicate in each trial, one individual was placed in the centre of the test arena underneath an upturned 70L nally bin for 10min to acclimate before being lifted for the trial to begin. To minimize disturbance, turtles were monitored via BluTooth GoPro video cameras attached and mounted to a suspended cross-bean overhanging each arena. Turtles were observed for up to 120mins to see if they could escape, after which the experiment ceased. After each trial, all turtles (including those that had escaped arenas) were kept in shaded, storage containers and released at the end of each day at the point of capture.

#### Experiment 2 – manipulated ‘gate’

We designed a second experiment to test whether turtles could locate ‘turtle gates’ if they could not fit through the standard pig meshing. All field arenas were set up with the small mesh on the bottom, as would be typical for a feral pig arena fence. An additional section of wire was weaved through the bottom row of wire meshing to ensure that turtles (44 *Emydura macquari krefftii*, and one *Myuchelys latisternum*) would not be able to pass through the fence without using the turtle gates (ensuring turtles were blocked in arenas – see Figure 1E). This permitted the use of a wide range in body sizes (even those that would normally be able to pass through the small meshing). Turtles were placed into arenas (described above) with ‘turtle gates’ clipped into the bottom row of the fence. We examined if and how long it took turtles to locate and successfully pass through gates using three distinct treatments: field arenas with gates every 1, 2 and 4m along the base. Each arena received the same gate spacing around the entire perimeter. The time it took turtles from release to exit through a gate after encountering a fence, and how far turtles travelled along the fence before existing the arena through a gate were recorded.

### 2.4 Data analysis

To examine whether turtle morphometrics differed between the Archer River floodplain (lower wetlands) to those captured in the upper catchment (upper wetlands), we used using multidimensional scaling ordinations, based on the Bray-Curtis similarities measure [44] with significance determined from 10,000 permutations. Multivariate dispersion were tested using PERMDISP, however, homogeneity of variance could not be stabilized with transformation, and therefore untransformed data were used. Multivariate differences using PERMANOVA [45] were tested using two factors: lower/upper wetlands (fixed), and fenced/unfenced (fixed).

## 3. Results

### 3.1 Archer River wetland field results

A total of 161 turtles were captured during this study, representing four species including *E. s. worrelli* (n=96), *Chelodina rugosa* (n=54), *M. latisternum* (n=6) and *C. canni* (n=5) (Table S3). There were 79 females, 63 males, 14 juveniles and 1 sub-adult captured (with four where sex could not be resolved). In addition, three individuals were identified from in situ shell material found adjacent to wetlands in both the upper and lower catchment. One *C. canni* and one *E. s. worrelli* were identified from in situ shell material found in the interior (not along the inside of the fence) of a fenced wetland in the upper catchment and one freshly pig predated, *C. rugosa* individual was found immediately adjacent to its aestivation site in an unfenced wetland located in the lower catchment.

The largest turtle captured was a female *C. rugosa* on the lower catchment floodplain, in an unfenced wetland (354.9mm SCL, 245.9mm SCW, 6.7kg wet weight), while the smallest was an *E. s. worrelli* in a fenced wetland in the upper catchment (95mm SCL, 87.5mm SCW, 110g wet weight). The average SCW (mean±SD) for each species was: *E. s. worrelli* (147.7±32.1mm, n=96)), followed by *C. rugosa* (160.7±33.5mm, n=54), *M. latisternum* (150.3±29.3mm, n=6), and *C. canni* (146.8±30.1mm, n=5).

There was an interaction between fencing/non fencing and wetland region in the catchment owing to a difference in the morphometrics for turtles between the lower and upper catchment wetland sites (PERMANOVA, interaction, Pseudo-F=5.81, *P*=0.02; Figure 2). However, some individuals from the unfenced lower catchment had turtles more similar to upper catchment fenced wetlands. Overall, turtles on the lower catchment floodplain were larger (including body weight) compared to those captured in the upper catchment.

**Fig 2.**
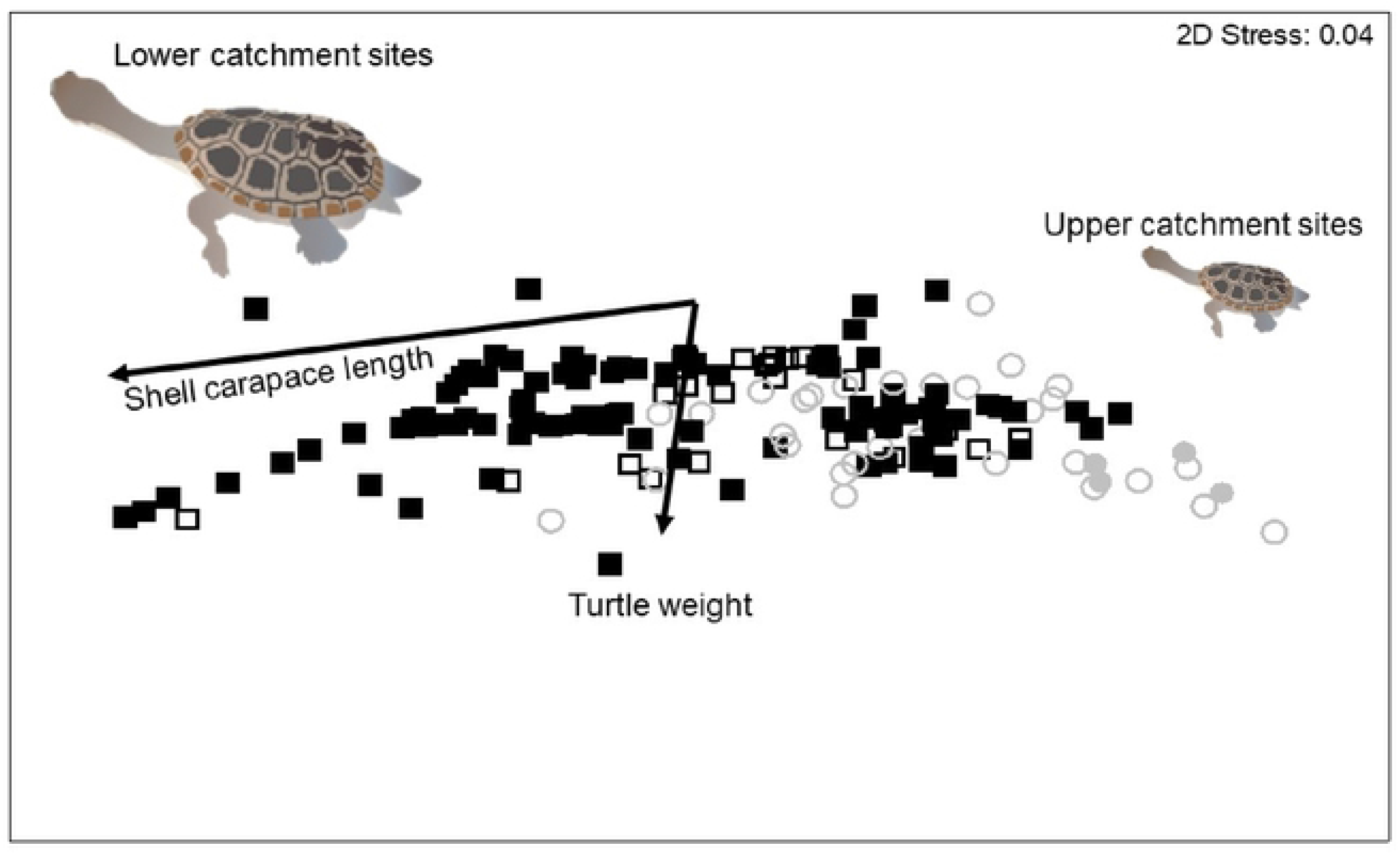
nMDS ordination of all individual turtles captured in the Archer River catchment during field surveys. Black boxes are turtles on the floodplain, grey circles are turtles from upper catchment - open symbols are fenced, and closed symbols are unfenced wetlands.

Pooling *C. rugosa*, *E. s. worrelli* and *M. latisternum* (161, 97% of total catch), 94 individuals caught in either fenced or unfenced wetlands that had a SCW greater than 150mm, and would likely not be able to negotiate exclusion fences. (It is possible that with the diagonal width of mesh approximately 180mm; Table S2, turtles with a SCW slightly greater than 150mm might squeeze through fence mesh though we could not confirm this at the time of field sampling and instead apply 150mm SCW to turtles – though see manipulative experiments below). Of the turtles captured, 44 individuals (46%) were captured inside fenced wetlands, predominately *E. s. worrelli* (32, 34%), and most caught in the upper catchment (Table 1), while the remaining 50 individuals were caught in unfenced wetlands in the lower catchment (*C. rugosa*). The remaining turtles (35) were smaller than 150mm and would be able to pass through fences.

**Table 1.**
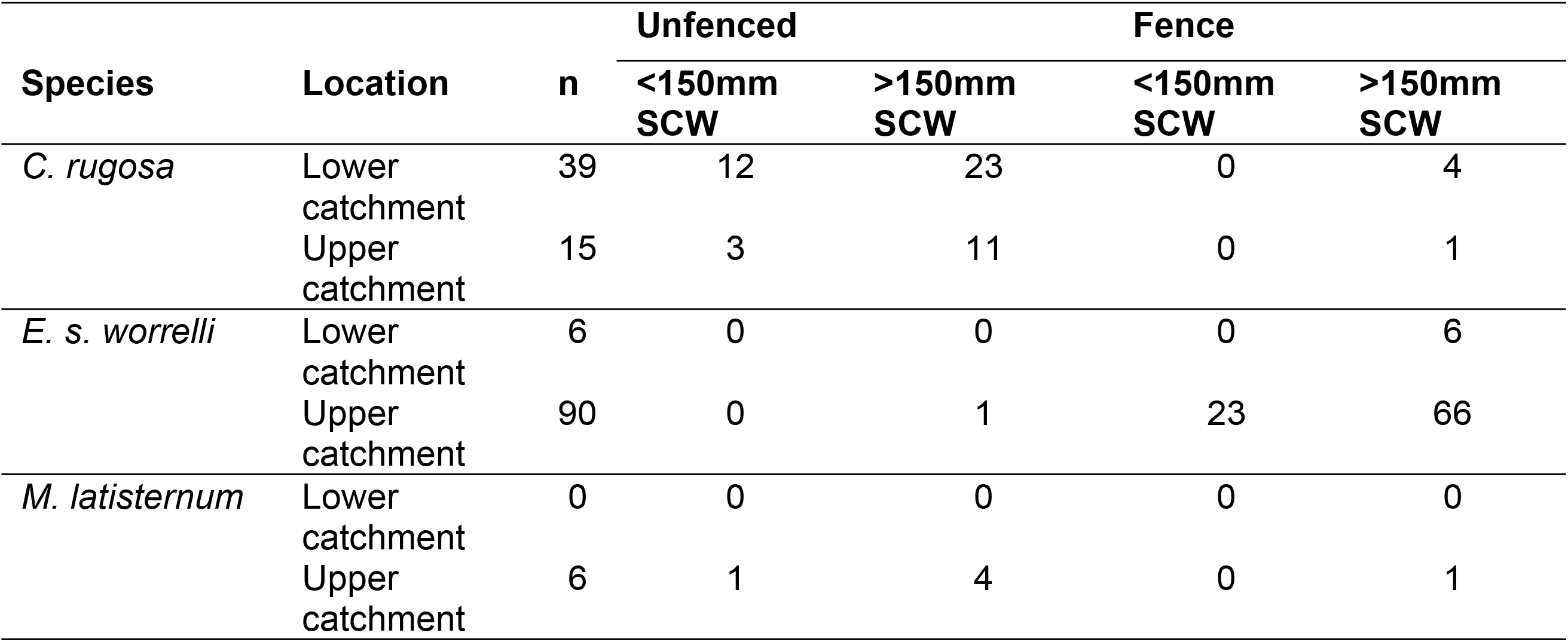
Summary of turtles captured in fenced and unfenced wetlands on the lower floodplain and upper catchment flood areas. *C. canni* not included here given turtles were found on road crossings, not in wetlands.

### 3.2 Fence manipulative experiments

#### Experiment 1 – mesh sizes

Sixty-five turtles (n=33 through small meshing; n=32 through large meshing) were used in this feral pig fencing experiment (Table 2). When deployed with the small size mesh closest to the ground, 78.6% (26/33) of turtles were able to pass through without becoming stuck. In contrast, nearly all turtles (98.6%; 31/32) were able to pass through the pig fences with the large square meshing on the bottom. Surprisingly, we also observed that even large turtles (with carapace widths wider than the meshing) were often able to escape through the fencing by lifting one side of their shell and passing through the mesh diagonally (Figure 1E). This is the first evidence to suggest that the primary limiting dimension of the fence meshing is the diagonal width, rather than a horizontal width, as suggested by the field data which was unable to indicate whether we could not say if those individuals would pass through fences or not.

**Table 2.** Size distribution of turtles from experiment 1 – passage rates through feral pig fencing. Turtles were either blocked or escaped (see Figure 1E). Fence mesh size represents the size mesh at the bottom of the fence, closest to the ground (large = 150×150mm; small = 150×100mm). SCW = straight carapace width; SCL = straight carapace length; carapace height = max height from plastron to carapace. Range represents minimum – maximum.

| Fence mesh size | Turtle outcome | n | Passage rate | SCW |  | SCL |  | Carapace height |  |
| --- | --- | --- | --- | --- | --- | --- | --- | --- | --- |
|  |  |  |  | Mean±SD (mm) | Range (mm) | Mean±SD (mm) | Range (mm) | Mean±SD (mm) | Range (mm) |
| Large | Blocked | 1 | 3.1% | 173.6 | 173.6 | 232.7 | 232.7 | 94.4 | 94.4 |
| Large | Escaped | 31 | 96.8% | 166.9 ± 15.0 | 139.5 - 205.8 | 218.3 ± 25.7 | 129.1 - 251.4 | 85.1 ± 10.1 | 59.7 - 101.0 |
| Small | Blocked | 7 | 21.2% | 177.6 ± 6.5 | 170.0 - 187.6 | 234.7 ± 6.5 | 226.0 - 245.0 | 94.4 ± 3.8 | 89.7 - 100.2 |
| Small | Escaped | 26 | 78.7% | 161.4 ± 13.9 | 121.4 - 184.5 | 210.8 ± 20.6 | 154.8 - 247.7 | 82.5 ± 9.5 | 63.2 - 100.0 |

#### Experiment 2 – installing gates

Turtles located gates after prospecting and trying to fit through meshing areas that were too small to pass through. The majority (92.1%, 35/38) of turtles was able to locate and pass through gates, regardless of their spacing, while 7.9% (3/38 turtles) failed to locate a gate within 2 hours (Table 3). For the three turtles that did not use gates, each appeared to have ceased attempts to pass through the mesh, dug into the grass, and remained motionless for the remainder of the trial. Gates applied every 4m showed an 83.3% passage rate (10/12 turtles), every 2m showed a 91.6% (11/12 turtles) passage rate, and turtle gates applied every 1 m showed a 100% passage rate (14/14 turtles). Turtles that used the gates spent less time searching for a passage through the fence when gates were closer together, with increased time searching with increasing distance between gates (Table 3).

**Table 3.** Passage rates of 38 turtles in experiment 2 – testing if turtles locate and use ‘turtle gates’. ‘Fence to escape’ represents the time turtles took to locate and use the turtle gate once they reached a fence. ‘Distance travelled’ represents the distance travelled once a turtle encountered a fence until it located a turtle gate, or the 2-hour time-cap elapsed.

| Turtle gate spacing (m) | Turtle outcome | n | Passage rate | Fence to escape (min) |  | Distance travelled (m) |  |
| --- | --- | --- | --- | --- | --- | --- | --- |
|  |  |  |  | Mean±SD | Range | Mean±SD | Range |
| 1 | Used gate | 14 | 100.0% | 3.7±8.5 | 0-33 | 2.0±1.5 | 0.1-4.6 |
| 1 | Blocked | 0 | 0.0% | - | - | - | - |
| 2 | Used gate | 11 | 91.6% | 6.3±12.7 | 0-43 | 1.9±2.2 | 0-6.3 |
| 2 | Blocked | 1 | 8.4% | 88 | 88 | 6.5 | 6.5 |
| 4 | Used gate | 10 | 83.3% | 8.8±10.6 | 0-36 | 2.1±1.6 | 0-4.5 |
| 4 | Blocked | 2 | 16.7% | 90.0±12.7 | 81-99 | 2.2±0.7 | 1.7-2.8 |

## 4. Discussion

Combing field and manipulative experiments, we reveal that most large turtles, which would not fit through existing pig fence designs, will prospect and move along a barrier fence until they find suitable passage. By applying gates every 1–4m can allow for nearly 100% passage rates of turtles that would otherwise be stuck on one side of the fence. Gates may be strategically applied in travel corridors [46] to minimize the need for large-scale clipping efforts around entire wetlands (see Figure 3) and would minimize the negative impacts on turtles by lowering energetic expenditure searching for a gate and reducing exposure to predation, overheating, and desiccation. Although untested, it is possible that installation of multiple gates may reduce the structural integrity of pig fences and result in breaches at weak points.

**Fig 3.**
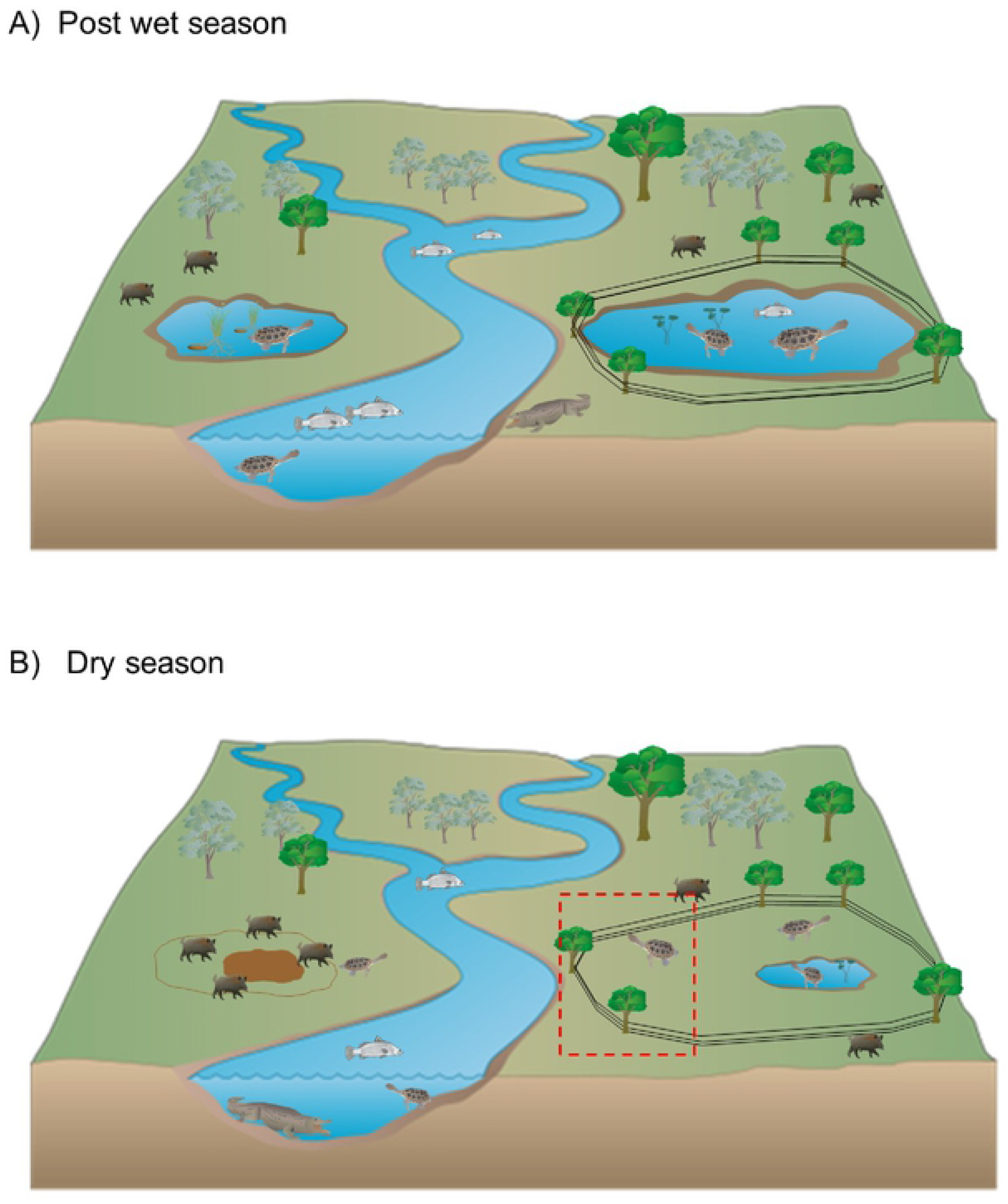
A) floodplain wetland complex following wet season and connection; B) floodplain late dry season with drying wetlands and impact of feral pigs, red dashed line illustrates where gates should be installed to maximise turtle escape and return to primary river.

While the installation of fences to exclude pigs from wetlands and the periodic culling of pigs remain common management strategies [22], our field study shows that fences can be detrimental for turtle populations. However this can be now overcome by incorporating gate modifications to fences to better assist freshwater turtles that have a shell width greater than the dimensions of the fencing wire would enhance their conservation. The data here shows that turtles, regardless of species, with a shell width greater than the diagonal wire gap will likely be trapped inside (or outside) fenced wetlands, limiting their access to important resources. The dilemma of reduced availability of freshwater turtle habitat can be mitigated by the simple and inexpensive design modification outline here, with turtles able to locate the gates and pass through them in a relatively short period.

Tropical wetlands can dry completely especially when they are not close to main river channels or permanent lagoons [26]. The rate of drying is dependent on antecedent wet season total rainfall, and the duration and frequency of floodplain connection [16]. Therefore in wet years the presence of water remaining in fenced wetlands is more likely after the onset of the wet season, which may for some species (Table S4) prohibit turtle overland dispersal to more permanent water. The wet season rainfall immediately prior, and during this survey, was within the 10^th^ percentile for historical records, which resulted in some wetlands drying out, requiring turtles to leave. In both cases, turtles are exposed to predation, either through pigs actively digging them up underground in unfenced wetlands (which was observed in this study), or during overland migration (by goannas, some bird species, wild dogs or pigs which are all predators of turtles).

Once erected, fence maintenance is imperative, particularly after bushfire, storm damage, or flooding that cause damage and compromise fences [47, 48]. Even after installing gates, surveys should continue to ensure that turtle movement throughout the landscape is not impeded by fences. Motion triggered cameras and passive transponder trackers [49] could be installed at gates while routine inspections along fences (as part of general maintenance) ensuring that gates are in the most effective location. Further modifications could be administered retrospectively after gates are installed.

The size separation in turtles between floodplain wetlands low in the catchment and riverine wetlands higher in the catchment was unexpected. This highlights important underlying differences in environmental conditions or food limitation contributing to turtle growth in the upper catchment remaining smaller compared to those on the expansive floodplain areas. This highlights the need to undertake extensive baseline surveys to understand local species morphology, as the inclusion of gate designs in wetland fences, even though inexpensive, might not be always necessary – which has the advantage of protecting fence integrity.

## 5. Conclusions

Each conservation fence program requires a scientific monitoring package to evaluate the efficacy, but more importantly to identify whether additional design improvements are necessary. We advocate here that an easy management response is to ensure the wider diagonal width squares are located along the ground when erecting fences, rather than the small diagonal width squares. This simple tactic increases the number of turtles that could pass through the fence without delay, and would conceivably not decrease the structural integrity of the fences to withstand pig prospecting. However, simply removing a small piece of wire to increase openings allows for nearly 100% passage rates of turtles that would otherwise be stuck on one side of the fence. Turtle gates may be strategically applied in travel corridors to minimize the need for large-scale clipping efforts around entire wetlands. Further, gates can be easily retrofitted to existing fence designs, which has enormous positive conservation benefits for turtles in an already challenging, and changing floodplain environment.

## Acknowledgements

This project builds on a long-term feral animal management and monitoring program developed by Kalan enterprises and Aak Puul Ngangtam (APN) and their partners. Kalan and APN have developed their feral animal research and management agenda to meet the objectives of traditional owners in the region and have invited science organisations (CSIRO, James Cook University and the Department of Science and Environment) to contribute to the outcomes. APN and Kalan have conducted systematic feral pig control and monitoring in the Archer River basin for the past 6yrs. This study is supported by the Australian Government National Environment Science Program (Northern Australian Hub) awarded to CSIRO, James Cook University, and the Queensland Government. Capture of turtles and gate exclusion experiments were conducted under JCU ethics approval A2359.

